# Elucidation of the molecular interaction network underlying full-length FUS conformational transitions and its phase separation using atomistic simulations

**DOI:** 10.1101/2025.04.28.651058

**Authors:** Shuo-Lin Weng, Priyesh Mohanty, Jeetain Mittal

## Abstract

Fused in Sarcoma (FUS), a multi-domain RNA-binding protein, orchestrates cellular functions through liquid-liquid phase separation (LLPS), which promotes the formation of biomolecular condensates *in vivo*. While crucial to understanding cellular processes, an atomic-level view of the interdomain interactions associated with full-length (FL) FUS LLPS remains challenging due to its low solubility *in vitro*. Here, using all-atom (AA) molecular dynamics (MD) simulations, we examined the conformational dynamics and interdomain interactions of FL FUS in both dilute and condensed phases. Comparing two modern force fields (FFs) - Amber ff03ws and ff99SBws-STQ, we found that monomer simulation ensembles generated by both FFs exhibited qualitatively similar intramolecular interaction profiles dominated by intrinsically disordered regions (IDRs). While the two folded domains minimally participated in interdomain interactions, their stabilities significantly influenced the chain dimension and led to discrepancies compared to experimental data for both FFs. We observed that the Amber ff99SBws-STQ coupled with bond parameters adopted from the Zinc Amber force field (ZAFF) maintained stable folded domains and improved estimates of the chain dimensions. Finally, a microsecond-timescale simulation of FL FUS condensate revealed an extensive network of electrostatic interactions which are strongly correlated with those that modulate the dilute phase chain dimensions. Overall, insights from our all-atom simulations illuminate the interplay between folded domain stability and IDR interactions in modulating protein conformation and phase separation, advancing our understanding of FUS-related pathologies at the molecular level and aiding in the development of new therapeutics.

## Introduction

Fused in Sarcoma (FUS) is a multifunctional RNA- and DNA-binding protein that comprises a disordered N-terminal low-complexity sequence (LC) domain, two folded domains – an RNA recognition motif (RRM), and a zinc finger (ZnF) domain, three arginine–glycine– glycine-rich (RGG) domains, and a short nuclear localization sequence (NLS) (**Fig. 1**).^1^ Collectively, RRM, ZnF, and RGG1-3 form the RNA-binding domains (RBDs). FUS has been demonstrated to undergo liquid-liquid phase separation (LLPS) *in vitro*,^2–5^ bind with nucleic acids,^6–8^ and participate in the formation of various membraneless organelles (MLOs) such as stress granules and nuclear bodies, contributing to cellular functions, including cellular stress responses, transcription, splicing, and DNA repair.^9–13^ Specifically, current studies indicate that FUS plays a critical role in several DNA damage response (DDR) pathways through its multivalency and ability to undergo LLPS, which are essential for initiating DDR and facilitating the recruitment and organization of repair factors at damage sites.^14–18^ Moreover, FUS is also shown to interact with RNA polymerase II to regulate the transcription process and gene expression.^19–21^ Notably, FUS dysregulation has been associated with neurodegenerative diseases and cancers, including Amyotrophic Lateral Sclerosis (ALS) and Frontotemporal Dementia (FTD), due to dysfunctional DNA repair, abnormal cytoplasmic mislocalization, and liquid-to-solid transition (LST) forming amyloid fibrils. The association establishes FUS as a key model for uncovering the molecular mechanisms underlying these pathologies.^3,22–26^

**Figure 1.**
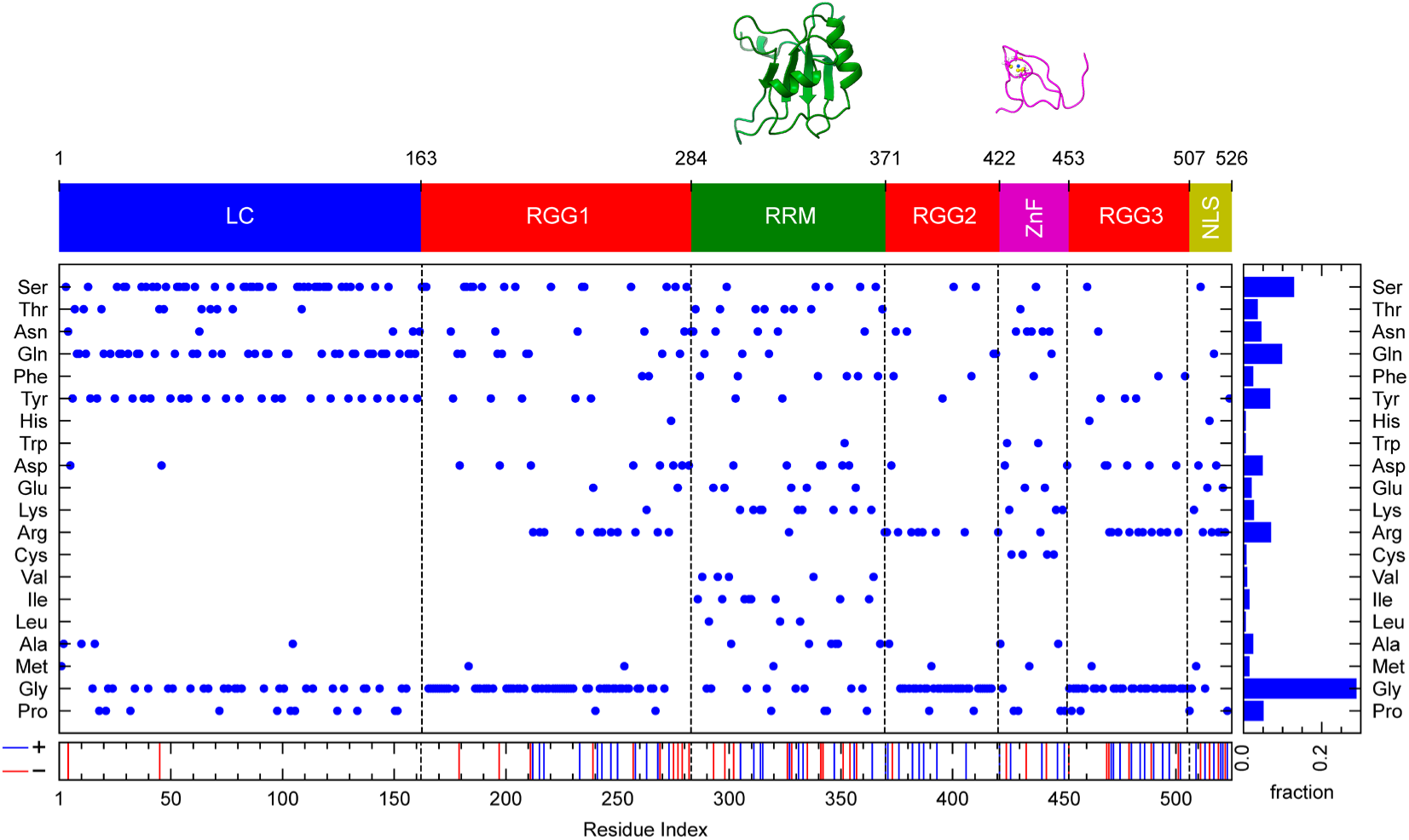
Domain structure and residue composition of FL FUS. FUS consists of a prion-like low-complexity domain (LC, aa 1-163), three arginine-glycine-glycine rich domains (RGG1, aa 164-284; RGG2, aa 372-422; RGG3, aa 453-507), two folded domains: the RNA-recognition motif-containing domain (RRM, aa 285-371) and the zinc-binding finger domain (ZnF, aa 423-453), and a nuclear localization sequence domain (NLS, aa 508-526). Residue composition is detailed for the FL sequence. The coloring scheme for each domain is consistent throughout the study. Structures of the folded domains are taken from the Protein Data Bank (PDB IDs: 6GBM and 6G99).

While the LC domain is capable of phase separation independently,^5,27–30^ RBDs significantly enhance and modulate LLPS.^2,20,31,32^ Previous mutagenesis experiments, solution-state nuclear magnetic resonance (NMR) studies, and computational investigations have demonstrated the major driving forces of FUS LLPS, including π-π, cation-π, hydrogen bonding, and electrostatic interactions.^5,20,27,31,33^ However, gaps remain in understanding the interplay between each domain in the context of full-length (FL) FUS, particularly the contribution of folded domains, due to the challenges associated with its low solubility.^2,25^ Specifically, there are insufficient atomistic details about the various types of interdomain interactions in FL FUS and a limited understanding of the role of folded domains in phase separation. Interestingly, the RRM domain of FUS and other aggregation-prone proteins exhibits low thermal stability (T_m_ ∼50 °C), which may promote misfolding and self-assembly into amyloid fibrils, crucial for FUS cytotoxicity *in vivo.*^34–36^ Stabilization of the RRM domain by the chaperone HspB8 can slow the aging process of FL FUS droplets, suggesting that RRM stability impacts intracondensate dynamics.^37^ On the other hand, NMR experiments hint at transient interdomain interactions between the RRM and other domains,^34^ although atomistic details are lacking. The ZnF domain adopts a folded conformation when bound to a zinc ion, and its removal can lead to reversible unfolding.^32,38^ The presence of excess zinc ions in solution can improve the stability of ZnF, enhancing the LLPS of FUS, and promoting the cytoplasmic migration of FUS.^38,39^ Understanding the molecular mechanisms underlying RRM misfolding, particularly the influence of interdomain interactions on its conformational stability, and on the overall chain dimensions and phase separation of FUS, can provide crucial insights into disease mechanisms and the development of targeted therapeutics.

To address the above-specified knowledge gaps, we utilized all-atom (AA) explicit-solvent molecular dynamics (MD) simulations to investigate the interdomain and residue-level interactions of FUS in the dilute (monomeric) and condensed phases. MD simulations, complementing experimental approaches,^40,41^ have been successfully applied to understand LLPS behavior of intrinsically disordered regions (IDRs) and multidomain proteins such as TDP-43, hnRNPA1, hnRNPA2, NDDX4, and FUS.^5,20,27–30,33,42–49^ By leveraging simulations at varying resolutions, these studies provide critical insights into conformational ensembles, molecular interaction networks, intra-condensate dynamics, and the structural evolution of biomolecular condensates. In this study, we first performed AA MD simulations of the FL FUS monomer using two modern force fields (FFs) - Amber ff03ws and ff99SBws-STQ,^50–52^ and characterized the conformational ensembles in terms of their overall chain dimensions and the underlying intramolecular (both interdomain and residue-level) interactions. Notably, the ff03ws ensemble displayed significant chain compaction with respect to experiment which was associated with the partial unfolding of both folded domains. In contrast, our analysis indicated that ff99SBws-STQ, incorporated with bond parameters adopted from the Zinc Amber force field (ZAFF),^53^ maintained the stability of both folded domains and provided a reasonable estimate of overall chain dimensions compared to experiment. With this modified FF, we performed a large-scale simulation of the FL FUS condensate for 2.5 microseconds, which revealed an elaborate network of electrostatic interactions and provided both a domain- and residue-level view of the interactions which stabilize the condensed phase. Further, our comparative analysis of the dilute and condensed phase contacts revealed a strong positive correlation for pairwise residue-level contacts formed in these phases, thereby enabling a logical extension of the established positive correlation between single-chain and condensed phase interactions of intrinsically disordered polypeptides (IDPs)^54–56^ to multidomain proteins. By elucidating how folded domains modulate FUS conformation and phase separation through structural fluctuations in addition to residue-level IDR interactions, our work advances the mechanistic understanding of FUS aggregation-related pathologies and provides a framework for developing condensate-specific therapeutics for neurodegenerative diseases.

## Methods

### All-atom MD simulations

#### Initial structures and molecular modeling

The initial conformations for AA simulations of the FL FUS single chain were taken from the HPS-Urry coarse-grained (CG) simulation with a single bead per-residue resolution.^57,58^ Based on the AlphaFold2-predicted structure (https://alphafold.ebi.ac.uk/entry/P35637),^59,60^ RRM (residues 285-369) and ZnF (residues 422-453) domains were set as rigid bodies, while the rest of the protein was kept flexible. The CG simulation was performed at 300 K using the LAMMPS software package.^61^ Random conformations were sampled over a 1-μs CG simulation and converted to AA configurations using the Modeller package,^62^ using solution-state NMR structures (PDB IDs: 6GBM and 6G99^63^) as templates for the folded domains.

For isolated folded domain simulations, protein structures were taken directly from the corresponding NMR structures.^63^ To model the FL FUS condensate, a configuration with 25 protein chains was extracted from a CG phase coexistence simulation conducted at 300 K using HOOMD-Blue 4.7.0.^64^ The Modeller package was used to reconstruct the all-atom slab configuration, again incorporating NMR structures as templates for the folded domains.^63^ This approach followed established protocols from our previous studies.^27,31,42^

#### Force field choice and simulation protocol

All systems were simulated in explicit solvent, modeled using the Amber ff03ws or ff99SBws-STQ force fields and the TIP4P/2005 water model,^50–52,65^ both incorporating improved NaCl parameters.^66^ Zinc ion binding within the ZnF domain was represented using two parameter sets: nonbonded (NBM) and bonded (ZBM) models. The NBM approach utilized CYZ parameters as outlined by Macchiagodena *et al.*,^67^ which we validated in a prior study examining zinc-mediated modulation of SOD1 protein conformations.^68^ The ZBM approach employed the Zinc Amber Force Field (ZAFF) developed by Peters *et al.*^53^ The protein molecules were placed in boxes of appropriate dimensions: an octahedron (edge length: 15 nm) for FL FUS single-chain simulations, a cubic box (edge length: 6 nm) for isolated folded domains, and a 12.5 × 12.5 × 50 nm tetragonal box for the FL FUS slab. The systems were solvated with counterions added to achieve electroneutrality and a salt concentration of 150 mM to mimic physiological conditions.

Each system underwent energy minimization with the steepest descent algorithm, followed by a 100 ps NVT equilibration at 300 K. For the condensate simulation, an initial 250-ps annealing cycle (5–300 K) with position restraints applied to backbone heavy atoms preceded the NVT equilibration. The velocity rescaling algorithm was used for temperature control with a coupling constant of 0.1 ps.^69^ Subsequently, a 100-ps NPT equilibration was conducted using the Parrinello-Rahman barostat with a coupling constant of 2 ps for pressure control at 1 bar.^70^ Initial equilibration simulations were performed using the classical MD package GROMACS-2022,^71,72^ employing periodic boundary conditions and a 2-fs integration timestep. Production simulations were performed using Amber 22 or OpenMM 8.1.1 for enhanced performance and flexibility.^73–75^ For Amber simulations, the structure and topology files obtained from GROMACS were converted using the ParmEd package in AmberTools23, while OpenMM simulations directly utilized the GROMACS files. Hydrogen mass repartitioning was applied in both platforms to enable a 4-fs timestep.^76^ All production simulations were performed in the *canonical ensemble*, with Langevin dynamics^77^ controlling the temperature at 300 K (friction coefficient = 1 ps⁻¹). Short-range nonbonded interactions were calculated with a 0.9-nm cutoff, and long-range electrostatic interactions were computed using the particle mesh Ewald (PME) algorithm.^78^ Bond constraints for hydrogen-containing bonds were enforced in OpenMM and Amber with the SHAKE algorithm. For the RRM domain, restraints to maintain the native folded state were implemented in OpenMM using the CustomBondForce class to apply a flat-bottom restraint potential based on NMR-derived hydrogen-bond information. The potential function is:

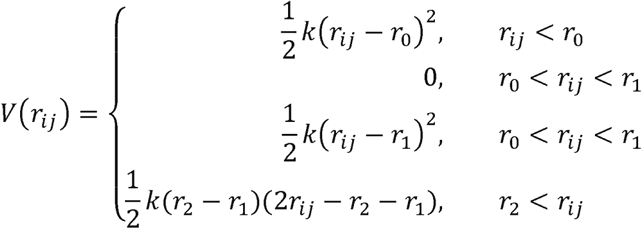

where k = 20 kcal/nm^2^, r_1_ = 0.27 nm, r_2_ = 0.3 nm, and r_3_ = 0.35 nm.

After production runs, the Amber NetCDF or CHARMM DCD trajectory files were converted to GROMACS compressed trajectory files via the CPPTRAJ package^79^ in AmberTools 23 for further analysis. A concise summary of the single-chain simulations is provided in **Supporting Table 1**. The system size of the all-atom slab simulation is described in **Supporting Table 2**.

### MD trajectory analysis

Protein-protein pairwise contacts were calculated using MDAnalysis 2.5.0,^80,81^ following the methodology defined in our previous work.^31^ A contact was considered formed if any heavy atom from two residues was within 4.5 Å. Residue pairwise contacts were determined by summing all contacts between heavy atom pairs from the respective residues, excluding interactions involving residues within a five-residue proximity. The *gmx polystat* command was used to compute the radius of gyration (R_g_) and end-to-end distance (D_ee_), while autocorrelation functions were calculated with the *gmx analyze* command with “*-ac*” flag. The hydrodynamic radius (R_h_) was calculated using our custom script implementing the HullRad algorithm^82^ with MDAnalysis,^80,81^ enabling trajectory-wide analysis with multiprocessing. Root-mean-square deviations (RMSD) and fluctuations (RMSF) for the folded domains were calculated with CPPTRAJ.^79^ Our established protocol was used to analyze angle-distance distributions,^31^ with the contact definition cutoff adjusted to 4.5 Å. Visualization of snapshots was performed using VMD or ChimeraX.^83,84^ The first 1 μs of each simulation was skipped as equilibration, based on the chain relaxation time estimated from the R_g_ autocorrelation functions in **Supporting information Figure 1** (**SI Fig. 1**). Density profiles of the condensed phase were generated from the final 200 ns of simulation trajectories using the *gmx density*. Local ion concentrations were predicted using a simple model proposed in prior studies.^27^ Except for the isolated ZnF simulation with ZBM, all statistical data from single-chain simulations were averaged over three independent trajectories. For the condensed phase trajectory, all data presented in this study were averaged over 25 chains.

## Results

### Conformational Ensembles of FL FUS Monomers Reveal Force Field-Dependent Chain Dimensions and Conserved Interaction Networks

We chose two state-of-the-art force fields to model the FL FUS monomer ensemble - (i) Amber ff99SBws-STQ^52^ and (ii) ff03ws.^50,51^ Both force fields have been previously shown by us and others to accurately describe the global chain dimensions and local structural features of a wide variety of IDPs and multidomain proteins.^27,31,42,85–87^ The radius of gyration (R_g_) and hydrodynamic radii (R_h_) are widely used metrics to characterize the overall chain dimensions and structural flexibility of multidomain proteins such as FUS, as most regions are disordered.^88^ Our simulations show distinct distributions of R_g_ between the two force fields, reflecting their differences in terms of the sampled conformations **(Fig. 2A, B)**. The ff03ws model produced a collapsed and less dynamic ensemble with a mean R_g_ of 2.6 ± 0.1 nm. In contrast, the ff99SBws-STQ model exhibited a more expanded conformational ensemble with a mean R_g_ of 5.0 ± 0.9 nm. Our results are similar to those recently reported by Sarthak *et al.*, where ff03ws gave rise to a collapsed ensemble while ff99SB-disp yielded expanded conformations at 150 mM NaCl.^89^ Sarthak *et al.* also reported an experimental R_g_ distribution ranging from 3–5 nm, measured by dynamic light scattering (DLS) experiments in the presence of 1 M KCl.^89^ However, such high salt concentrations are likely to impact the conformational ensemble of FUS due to charge screening or salting-out effects.^31,90^ Likewise, Yoshizawa *et al.* reported an expanded conformational ensemble (R_g_ = 4.87 nm) for maltose binding protein (MBP)-tagged FUS in the presence of a moderate high salt concentration (0.5 M NaCl) from small-angle X-ray scattering (SAXS) experiments.^91^

**Figure 2.**
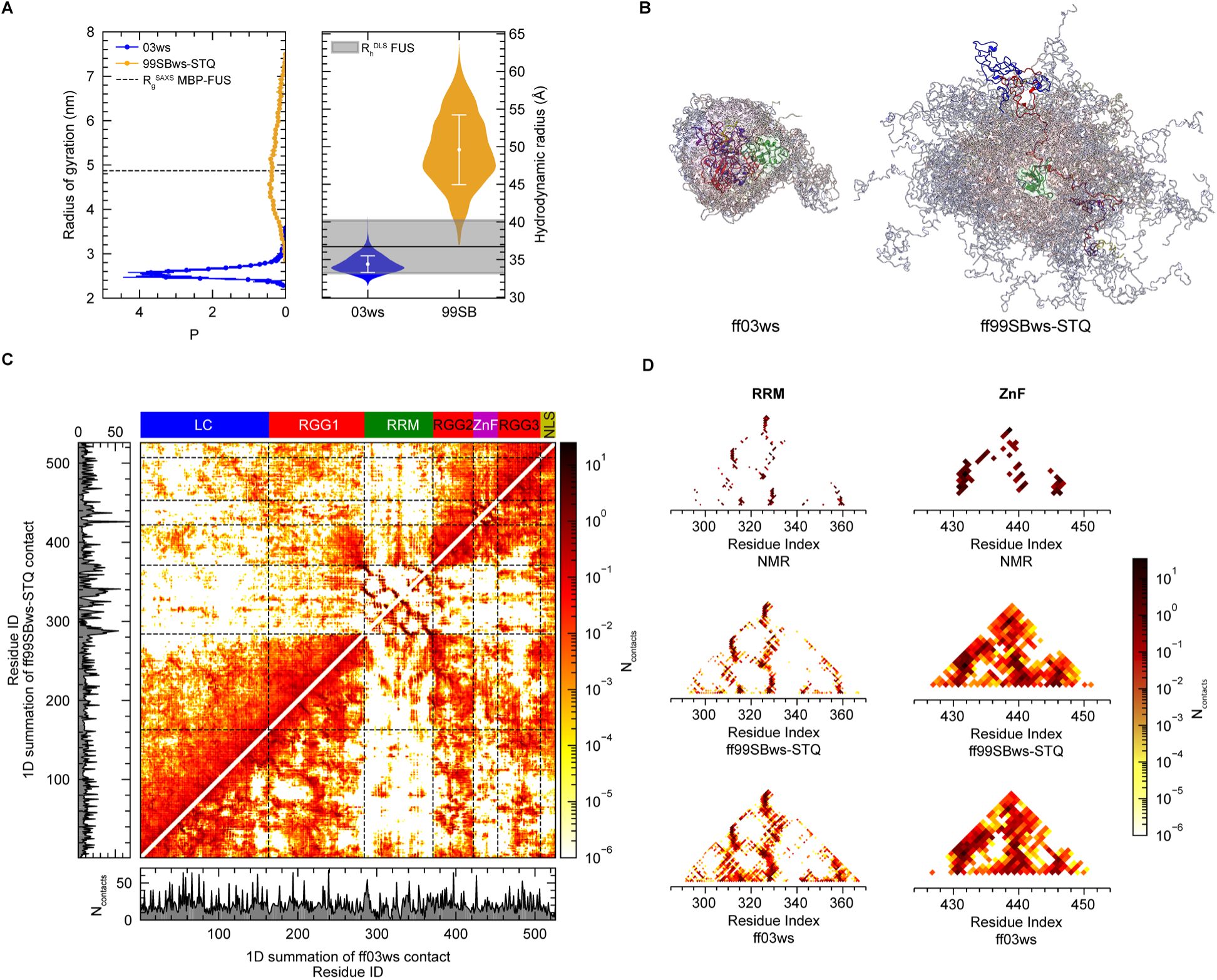
Conformational ensembles and interaction profiles of the FL FUS monomer. **A.** Left: Distribution of the radius of gyration (Rg) from atomistic simulations using ff03ws and ff99SBws-STQ force fields, along with the experimental MBP-FUS value. Right: Violin plot of hydrodynamic radius (Rh) distributions from both models and the experimental value. **B.** Snapshots of conformational ensembles of FL FUS. **C.** Intramolecular contact profiles showing heavy atom contacts for each position, averaged over simulation ensembles. The left column and bottom row represent one-dimensional summations. The top color bar indicates domain ranges. **D.** Contact profiles of the folded domains (RRM and ZnF) from simulation and NMR ensembles.

Under a moderate salt concentration of 200 mM NaCl, Sukhanova *et al.* reported the average hydrodynamic radius (R_h_) of FUS from DLS experiments as 3.67 ± 0.35 nm.^17^ For comparison, the R_h_ calculated from the ff03ws simulations using the HullRad algorithm^82^ was 3.44 ± 0.12 nm, while the ff99SBws-STQ simulations yielded an R_h_ of 4.96 ± 0.46 nm **(Fig. 2A)**. These results indicate that the mean R_h_ of FL FUS lies much closer to ff03ws although the overall chain dimension is still underestimated by ∼7%. Overall, while each model captures aspects of the experimental conformations, neither fully encompasses the experimentally observed native state ensemble, reflecting the inherent limitations of these force fields in fully reproducing FUS’s conformational diversity and challenges associated with experimental determination of R_g_/R_h_.

To explore the relationship between the observed chain dimensions and interdomain interactions in the FL FUS conformational ensembles, we computed the intramolecular contacts for both force fields as shown in **Fig. 2C**. Previous studies of IDPs suggest that the intramolecular interactions governing single-chain conformations in the dilute phase largely resemble those promoting LLPS for low complexity IDR sequences such as those occurring in FUS.^31,44,54–56,87^ Additionally, for multidomain proteins, specific intra- and interdomain interactions also affect their propensity to undergo LLPS.^42,92,93^ Interestingly, despite the quantitative differences in terms of overall contact counts due to distinct conformational preferences, both models exhibited qualitatively similar contact patterns (**Fig. 2C**). Extensive contacts were observed within the LC domain and between the LC and RGG domains, consistent with previous studies.^20,31,32,34^ Overall, our analysis reinforces the critical roles of LC intradomain and LC-RGGs interdomain interactions in FUS LLPS. Interestingly, upon closer inspection of the contact maps for the folded domains (**Fig. 2D**), we observed both RRM and ZnF domains in the ff03ws ensemble displayed a significant disruption of native pairwise contacts compared to the NMR structures, along with the emergence of numerous non-native contacts. In case of ff99SBws-STQ, while the RRM domain appears to maintain its native contact profile, non-native contacts were observed in the ZnF domain.

In conclusion, our simulations revealed distinct conformational ensembles for FL FUS monomers across the two force fields, yet they exhibited qualitatively similar interaction profiles. These profiles were characterized by extensive contacts within IDRs and limited contributions from folded domains, highlighting the central role of IDR interactions in driving LLPS and the contributions from all residues.^5,20,31^ Next, given that two force fields exhibited significant differences in contact formation for both folded domains, we aimed to determine whether these differences could explain the differences in the conformational ensembles generated using both force fields. To this end, we examined their stabilities in the FL FUS construct based on established structural metrics and compared them to those of their isolated counterparts.

### Folded domain stability modulates the conformational ensemble of FL FUS monomer

While *in vivo* experiments have linked the stability of the RRM domain to the aging of FUS droplets, the molecular details and mechanisms remain unclear.^32,34,37,38^ The RBD segment cannot phase separate on its own but can undergo LLPS when interacting with nucleic acids such as RNA, DNA, and ATP, and has been shown to enhance the LLPS of the LC domain.^8,20,32,94^ However, prior studies often included contributions from other domains, such as the RGG regions, leaving the specific role of folded domains insufficiently understood[NO_PRINTED_FORM]. As stated previously, comparisons with native NMR structures reveal significant disruption of intradomain contacts in the RRM/ZnF domain with ff03ws and the ZnF domain with ff99SBws-STQ (**Fig. 2D)**, implying destabilization of these domains in both force field ensembles. To further assess the stability of folded domains within the FL FUS ensembles, we computed the root-mean-square deviation (RMSD) and fluctuation (RMSF) of their Cα atoms, using the experimental structures as references (PDB IDs: 6GBM for RRM, 6G99 for ZnF).^63^ We also compared these results to those of isolated folded domain simulations to distinguish the influence of the force fields from domain interactions within FL FUS.

The RRM domain of FUS shares a common structure with RNA-binding proteins like TDP-43, hnRNPs, and the FET family, featuring a domain architecture with a central β-sheet flanked by two α-helices (**Fig. 1**).^63,95^ This domain has been shown to spontaneously self-assemble into cross-β-rich amyloid fibrils.^34^ The FL droplet with chaperone-stabilized RRM or RRM deletion displayed a delayed aging process.^37^ Additionally, structural comparisons between the isolated and the RNA-bound RRM indicate that RNA binding does not impact the RRM structure significantly.^63,95^ The ff03ws ensemble was found to populate high RMSD values exceeding 4 Å for RRM, while the ff99SBws-STQ ensemble exhibited a relatively stable RRM (**Fig. 3A, B** and **SI Fig. 2A**). Similar to our simulations, Sarthak *et al.* also reported that the ff03ws ensemble was characterized by an unstable RRM domain.^89^ Overall, these observations are also consistent with previous reports of folded state destabilization for ff03ws,^96,97^ which likely arises due to an overestimation of protein-water interactions. Interestingly, RRM stability was lower in the FL context compared to the isolated form for both force fields (**Fig. 3A**). RMSF analysis indicated that the increase in RMSD in the full-length protein was associated with a global increase in the conformational flexibility of the domain backbone for both force fields (**SI Fig. 2D**). Overall, the difference in stability between full-length and isolated domains suggests that the presence of disordered domains in the context of FL FUS destabilizes the RRM domain and increases its conformational flexibility.

**Figure 3.**
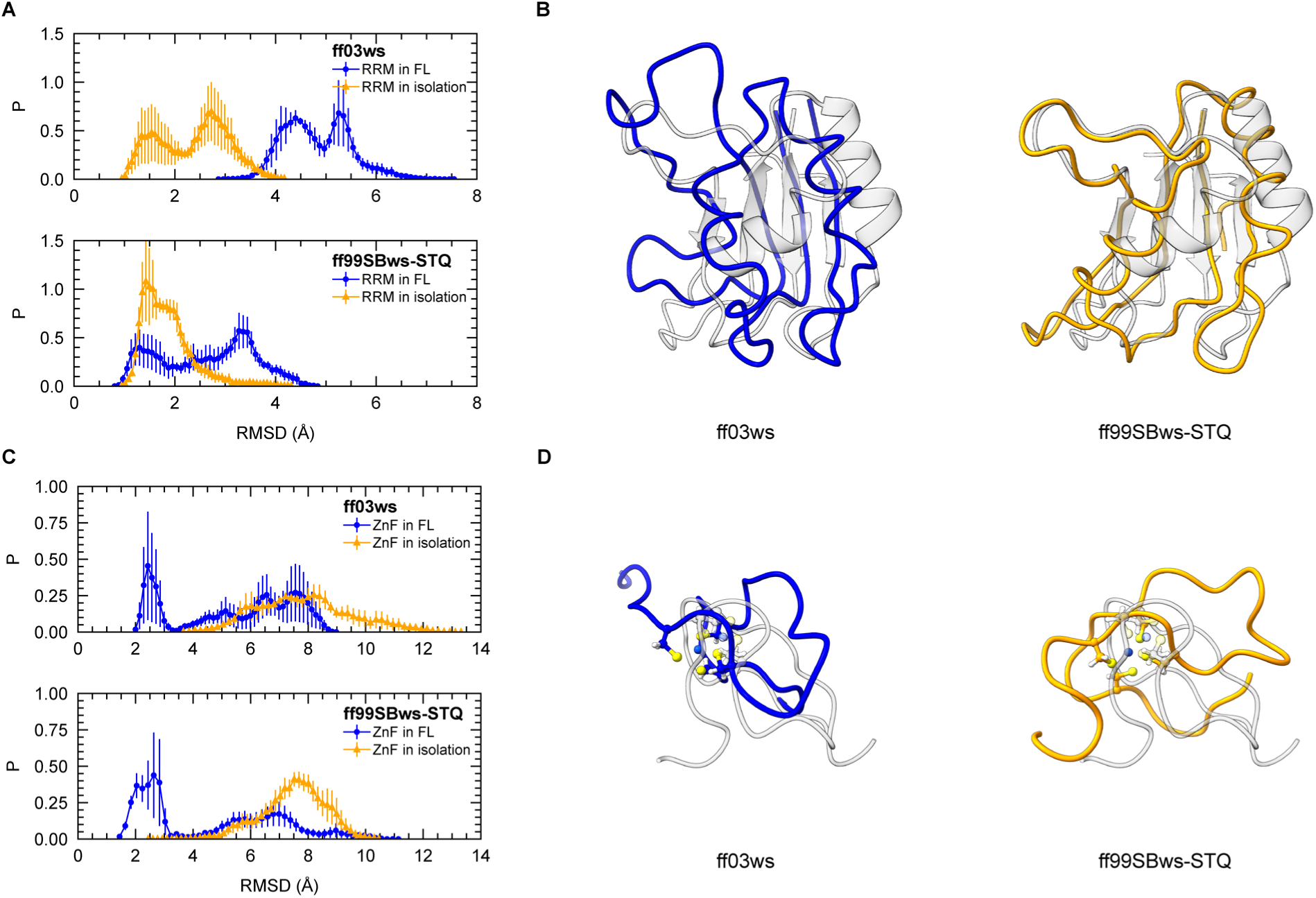
Comparison of folded domain stabilities between isolated and FL FUS constructs. **A.** Distribution of average root-mean-square deviation (RMSD) of Cα atoms in the RRM domain (residues 285-369) from three independent replicas for each condition. RMSD is calculated relative to the reference PDB structure (see Figure 1). **B.** Structure alignments to the 6GBM NMR structure (gray) from FL simulations. **C.** Distribution of average RMSD of Cα atoms in the ZnF domain (residues 423-453) from three independent replicas for each condition, relative to the reference PDB structure (see Figure 1). **D.** Structure alignments to the 6G99 NMR structure (gray) from FL simulations.

After investigating the RRM domain, we then examined the ZnF domain, which is part of the RanBP2-type ZnF family and features a typical four-cysteine motif that binds to zinc. When a zinc ion and single-stranded RNA containing the GGU motif are present, the ZnF domain forms a structure with two crossed β-hairpins (**Fig. 1**).^63^ The removal of zinc causes the ZnF domain to unfold (**SI Fig. 2C**), but this unfolding is reversible upon reintroducing zinc ions.^32^ In order to maintain stable coordination of zinc to the cysteine tetrad in ZnF, we utilized nonbonded Zn-Cys interaction parameters.^67^ The simulations with non-bonded parameters showed partial to complete unfolding of ZnF in both force fields with the presence of zinc, whether in isolation or in the FL context (**Fig. 3C, D** and **SI Fig. 2B**). Interestingly, unlike RRM, the ZnF domain is more stable and less flexible in the FL context compared to the isolated form (**Fig. 3C** and **SI Fig. 2E**), suggesting that the presence of other domains might enhance the conformational rigidity of ZnF. As with RRM, the ff99SBws-STQ model exhibits overall higher stability than ff03ws. Importantly, throughout most simulations, zinc remained bound to the cysteine residues, with small zinc-sulfur distances (**SI Fig. 3A**). Only one trajectory for isolated ZnF simulations showed dissociation for one cysteine, occurring after unfolding (**SI Fig. 3B, C**). Therefore, ZnF unfolding was not driven by the complete dissociation of zinc from ZnF. In conclusion, these unfolding results suggest the inability of nonbonded Zn-Cys interaction parameters to fully maintain the ZnF folded state, and stronger zinc binding via covalent bonds might be critical to preserving the ZnF native structure. Therefore, we tested the ability of Zn-Cys bonded interaction parameters to maintain the native structure in the next section.

### Stabilization of folded domains leads to convergence of single-chain MD ensembles between both force fields

To better understand the influence of folded domain stability on the overall conformation of FL FUS, we implemented specific modifications aimed at improving domain folding. For the RRM domain in the ff03ws model, we applied additional distance restraints derived from NMR structures to maintain its secondary structure (see Methods), effectively limiting its RMSD to less than 3 Å (**SI Fig. 4A**). These restraints resulted in a significantly expanded overall protein conformation compared to the original ff03ws model (**Fig. 4A**). This expansion likely arises because residues previously exposed in the unstable RRM, which interacted with other domains and promoted collapsed conformations, are no longer accessible. As anticipated, the RRM restraints altered the interaction profile, reducing RRM interdomain contacts and concentrating these interactions on specific surface-exposed residues. Additionally, the contacts between the LC and RGG2/3 domains were notably reduced, likely due to the decreased flexibility of the RRM caused by the restraints (**SI Fig. 4B**). These findings demonstrate that stabilizing the RRM domain affects the overall conformation and interdomain interactions within FL FUS.

**Figure 4.**
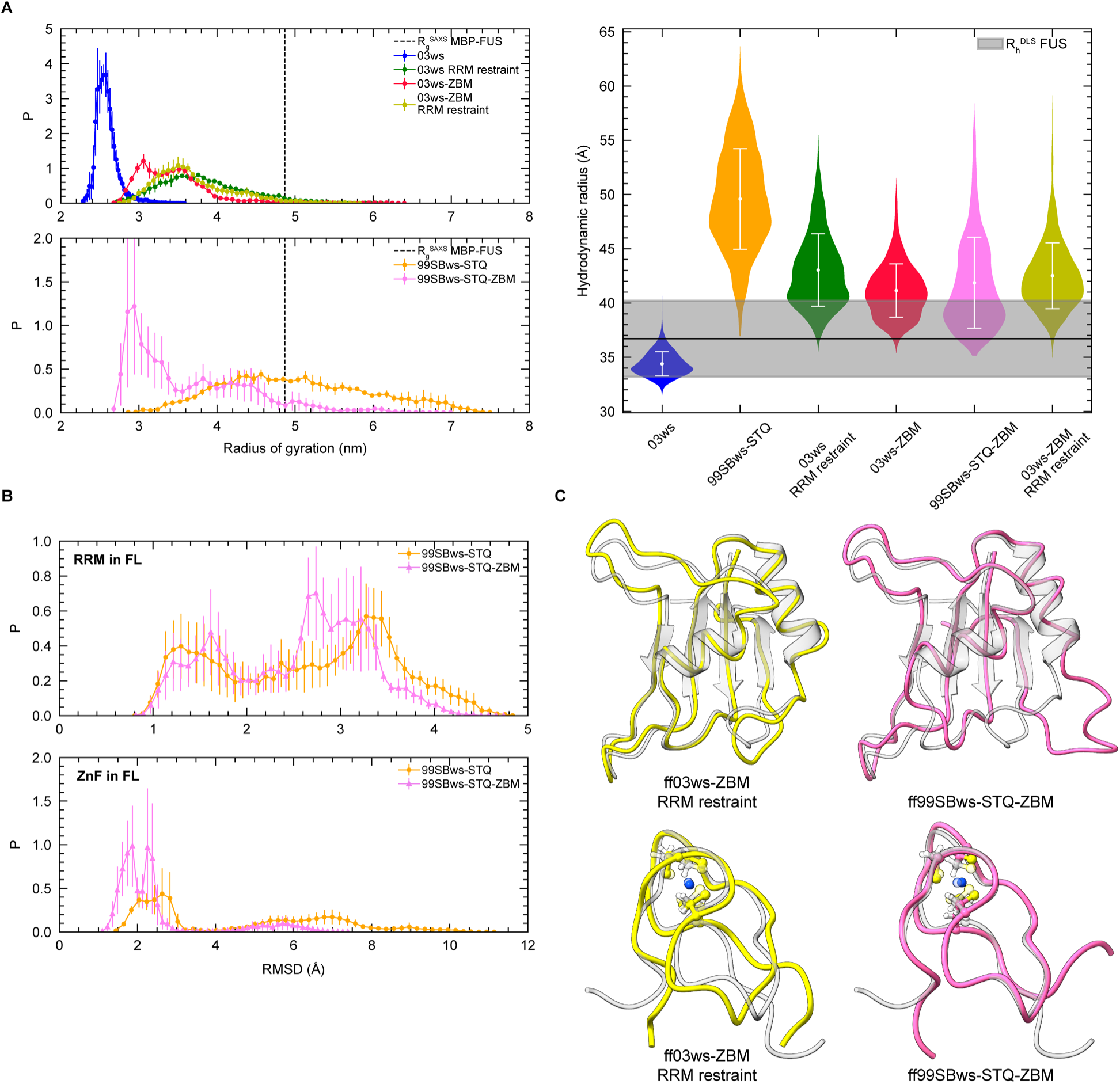
Increased stability of folded domains alters the chain dimensions of the FUS single-chain ensemble. **A.** Left: Distribution of Rg from atomistic simulations of FL FUS with different force fields, along with the experimental MBP-FUS value. Right: Violin plot of hydrodynamic radius (Rh)distributions for each force field and the experimental value. **B.** Distributions of average RMSD of Cα atoms in the folded domains from two ff99SBws-STQ force fields. **C.** Structure alignments to the NMR structures (gray) from FL simulations.

To improve the folding of the ZnF domain, we incorporated the well-established Zn-Cys bonded parameters from the Zinc Amber force field (ZAFF).^53^ The RMSD analysis indicated that the ZAFF parameters significantly improve the stability of ZnF native structure in both force fields, with RMSD values less than 3 Å for both the connected and isolated forms (**SI Fig. 4C**). These results are in line with the observations made by Sarthak *et al.* wherein force fields using covalent bonds to describe Zn-Cys interactions showed higher ZnF stability.^89^ Analysis of Zn-S distances showed that the ZAFF parameters resulted in longer Zn-S bond lengths compared to the nonbonded model (NBM), indicating that stabilization arises not from tight binding but from improved structural constraints (**SI Fig. 4D**). Additionally, Zn-S-Cβ angle analysis revealed a narrower range of distribution in the zinc bonded model (ZBM), implying that angle constraints contributed to improved ZnF folding (**SI Fig. 4E**). However, these bond lengths remained longer than those derived from the NMR structure,^63^ and the angles of the four cysteines were the same in simulations, while the angles from the NMR structure varied, suggesting that further refinement of the ZBM may be required to improve the overall agreement with experimental results.

Interestingly, the ZBM slightly compressed the protein chain for the ff99SBws-STQ ensemble, while causing chain expansion for the ff03ws ensemble. Applying both RRM restraints and the ZBM in the ff03ws model resulted in a further expanded conformation (**Fig. 4A**). Moving forward, we identified the ff99SBws-STQ force field with the ZBM as the most suitable model for describing FL FUS behavior, providing a reasonable estimate of chain dimensions with respect to experiment (both R_g_ and R_h_) (**Fig. 4A**), acceptable RRM stability, and significantly improved ZnF folding (**Fig. 4B, C**). Compared with the nonbonded model, their interaction profiles are similar, displaying dominant LC and RGG interactions with limited interdomain interactions involving folded domains. In the ZnF region, contacts are more concentrated and interdomain contacts are reduced, similar to the difference in the RRM region between ff03ws with and without RRM restraints (**SI Fig. 4E**). Overall, these results highlight that while folded domains in their native state contribute minimally to direct interactions within FL FUS, their unfolding can profoundly impact the conformational ensemble and intramolecular interaction profile. These simulations underscore the importance of accurate modeling for folded domains in understanding the conformational properties of FUS and other multidomain proteins.

Interestingly, a strong linear correlation emerged between the pairwise summations of contacts for both force fields when structural restraints are applied to stabilize folded domains (**Fig. 5** and **SI Fig. 5**). This indicates that contact-prone residue pairs are similar, suggesting both models captured comparable intramolecular interaction contributors. Arginine, tyrosine, glutamine, serine, and glycine residues emerged as dominant contributors to intramolecular contacts across all simulations. Beyond the traditionally recognized major contributors to LLPS – tyrosine-tyrosine and arginine-tyrosine pairs – significant contributions from polar residues like glutamine and serine were also observed.^2,31,87^ Notably, additional pairs involving arginine with other residues, particularly arginine-arginine interactions via π-π stacking, were identified, consistent with previous NMR and MD studies.^31^

**Figure 5.**
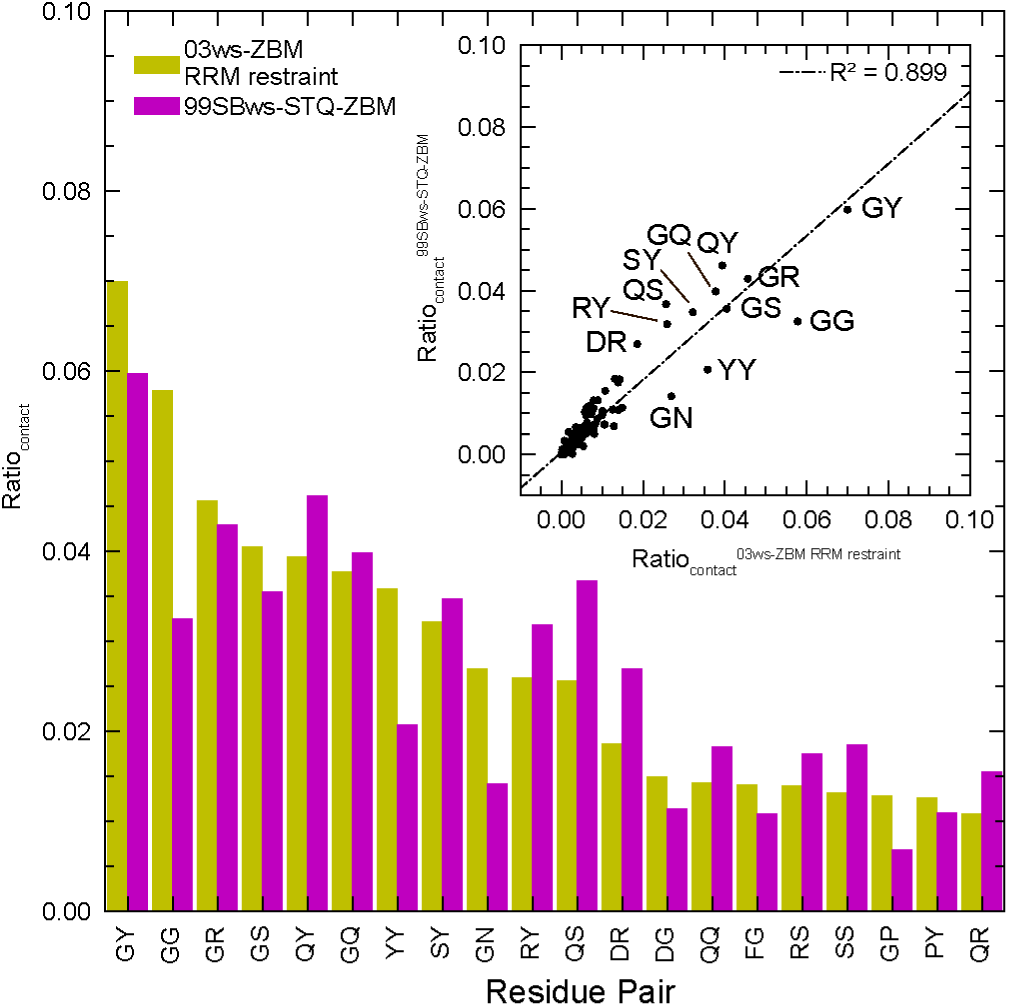
Residue-type interaction pairs modulating the monomeric conformational ensemble of FL FUS. The top 20 residue-pair contacts from FL FUS single-chain simulations with conformational restraints applied to RRM/ZnF domains are shown. See Supporting Figure 5 for full rank. Inset: Correlation between pairwise contact ratios in the ff03ws-ZBM RRM restraint (x-axis) and ff99SBws-STQ-ZBM (y-axis) models.

### FL FUS condensate simulation reveals an elaborate network of electrostatic interactions with enhanced folded domain stability

After investigating the interplay at the single-chain level, we aimed to characterize the network of intermolecular interactions and stabilities of the folded domain within the crowded environment of the condensate. To this end, we performed a large-scale simulation of the FL FUS condensed phase using a slab configuration containing 25 protein chains with the ff99SBws-STQ-ZBM force field (**Fig. 6A**), starting from an equilibrated state of the coarse-grained (CG) phase coexistence simulation. With the availability of high-resolution atomistic simulation data, we examined water and ion partitioning within the condensed phase (**Fig. 6B**). The FL FUS condensate protein concentration was approximately 500 mg/mL, with a water content of about 600 mg/mL, similar to previous FUS LC condensate simulations.^27,30^ Due to net positive charge of the FUS FL sequence, Cl^-^ ions exhibited a higher population inside the condensate compared to Na^+^ ions to maintain electroneutrality.

**Figure 6.**
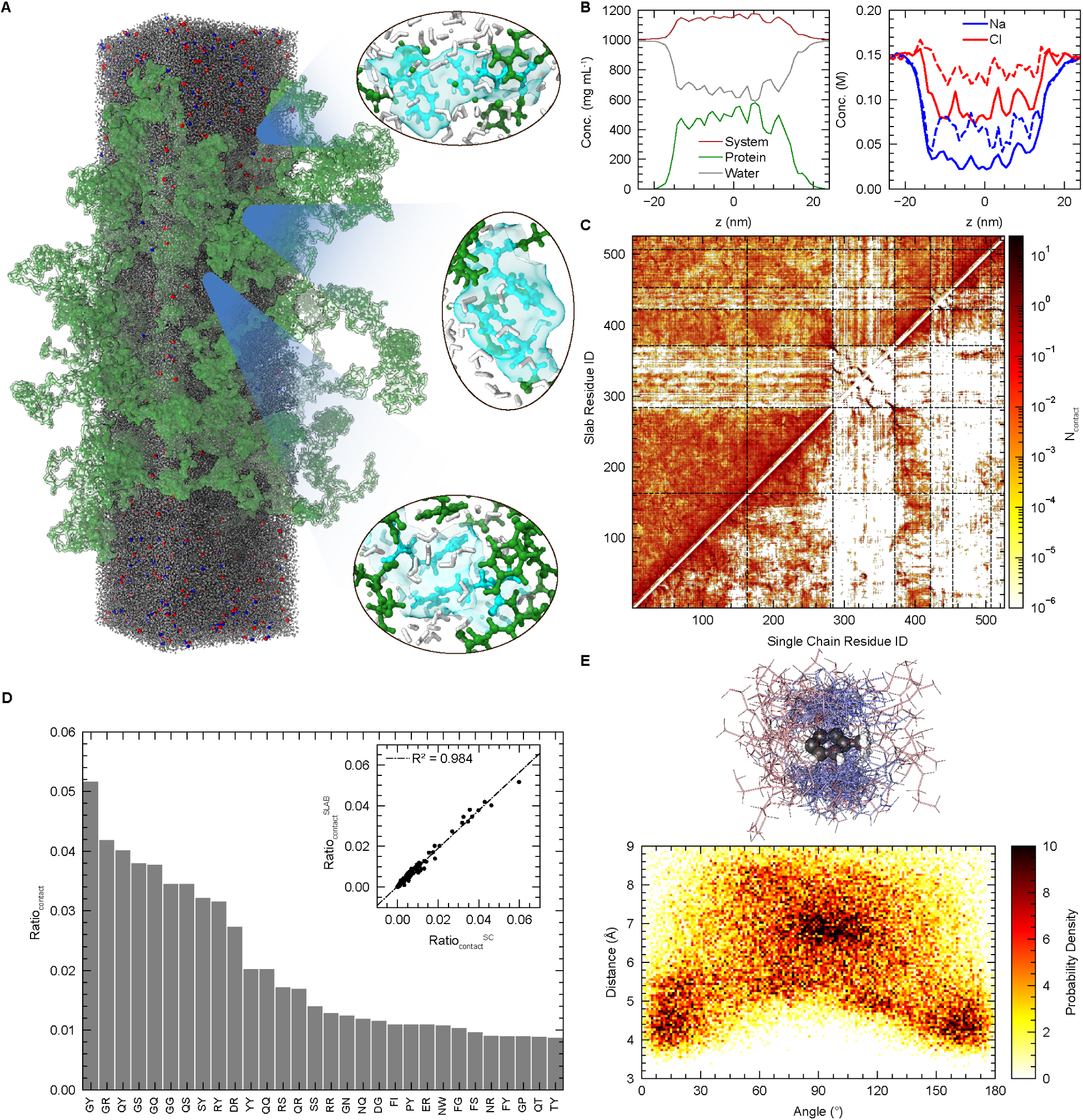
Single-chain interactions mirror those in FL FUS condensates. **A.** Atomistic condensed phase simulation of 25 chains of FL FUS (green), with water (gray), Na (blue), and Cl (red). Zoomed image of representative pair interactions and surrounding atoms. **B.** Density profiles from atomistic FL FUS condensed phase simulation. Components are shown in the legend. Dashed lines in right indicate the predicted ion concentrations by using concentrations of protein cationic and anionic residues. **C.** Contact profiles comparing FL FUS single-chain and slab simulations using ff99SBws-STQ-ZBM force field. Single-chain data averaged from three independent trajectories; slab data averaged across 25 chains. **D.** Top 20 residue pairs with highest contact frequencies in FL FUS slab simulation. Inset: Correlation of pairwise contact ratios between single-chain (x-axis) and slab (y-axis) simulations. **E.** Top: Representative configurations illustrating F-R contacts in FL FUS slab simulation. Bottom: Angle-distance distribution of F-R contacts.

Comprehensive pairwise contact mapping reveals universal residue participation in LLPS (**Fig. 6D** and **SI Fig. 6D**), with tyrosine, arginine, glutamine, serine, and glycine forming dominant contacts – consistent with their established roles in IDP phase separation and mirroring observations in FUS LC-RGG1 and other IDP condensates.^2,31,87^ Electrostatic interactions substantially contribute to the FL FUS interaction network due to abundant charged residues,^2^ while folded-domain hydrophobic residues show limited LLPS involvement despite forming stable intramolecular contacts as evidenced by pairwise two-dimensional contact mapping (**Fig. 6C**). Notably, we identified previously hypothesized but unverified phenylalanine-arginine (FR) interactions in FUS involving π-π stacking, confirmed through sp²/π plane angle-distance distribution analysis (**Fig. 6E**).^31,98,99^ This multimodal interaction landscape – combining stereospecific strong contacts with abundant weak interactions – demonstrates how diverse physicochemical forces synergistically mediate phase separation in multidomain systems.^31,87^

Consistent with observations in FUS LC, LC-RGG1, and other IDP condensates, the protein chains expanded in the condensed phase.^27,31,100^ The distribution of R_g_ and D_ee_ shifted towards larger values in the slab simulation compared to the single-chain simulations (**SI Fig. 6A**), resulting from a shift from intramolecular to intermolecular interactions within the condensate. Importantly, both RRM and ZnF domains remained folded for all chains throughout the trajectory (**SI Fig. 6B**), indicating that the condensate environment did not significantly destabilize these structured regions. Previous studies on IDPs have highlighted the relationship between single-chain properties and condensate behaviors, suggesting that intramolecular interactions observed in the dilute phase can contribute to condensate stability.^27,31,42,54–56,92,101^ Analysis of the contact maps within the FL FUS condensate revealed similarities and differences compared to the single-chain profile (**Fig. 6C**). As expected, interdomain contacts of the folded domains increased within the condensate due to enhanced opportunities for intermolecular interactions. However, the overall contact profile in the condensate, encompassing both inter- and intramolecular interactions, showed remarkable similarity to the single-chain ensemble, underscoring the dominant role of IDR interactions and the limited contributions from folded domains. This similarity is further supported by the high correlations observed in the per-residue and pairwise contact profiles (**Fig. 6D** and **SI Fig. 6C**), suggesting that the interactions governing monomeric conformations also play a crucial role in promoting and stabilizing LLPS,^54–56^ not only in IDRs but also in multidomain proteins.

## Discussion

FUS is a pivotal protein in various cellular functions due to its multivalency and capability to undergo LLPS. Understanding the mechanisms behind its self-assembly and the interactions between its domains is essential for elucidating its roles in cellular processes. FL FUS contains IDRs and structured domains such as the RRM and ZnF, which together contribute to its dynamic behavior and interactions.^1^ Investigating the interplay between these domains enhances our comprehension of FUS phase behavior and its implications in cellular organization and disease-related aggregation. Our study employed AA MD simulations to investigate the FL FUS interaction network, utilizing two different force fields with nonbonded zinc parameters. Nonbonded zinc models highlighted extensive contacts dominated by aromatic, arginine, and polar residues within IDRs, with minimal contributions from folded domains. These findings reinforce that FUS phase separation is primarily driven by IDRs, with contributions from all residue types. While tyrosine and arginine have traditionally emphasized roles, our results highlight underappreciated contributions of glutamine, serine, and glycine to the collective interaction network.^31,87,102^

Beyond interactions, our results illuminate the significant influence of folded domains on the overall conformational ensemble and phase behavior of FL FUS. Simulations of isolated folded domains demonstrated that the RRM domain is more stable in isolation than within the FL context, suggesting that interdomain interactions destabilize RRM. Specifically, the ff03ws model, with an unstable RRM domain, exhibited a collapsed conformational ensemble, while the same model with a stabilized RRM domain by external restraint had a more expanded protein conformation. This relationship between RRM stability and global conformation provides a mechanistic basis for the observed rapid aggregation of wild-type FL FUS droplets and the slower aggregation with stabilized RRM, emphasizing the critical role of RRM stability in modulating FUS phase dynamics. Similarly, our simulations revealed that bonded zinc-coordination parameters are essential for maintaining the folded state of the ZnF domain. The introduction of ZBM significantly improved ZnF domain stability and folding accuracy, providing insights into the structural preservation within FUS and guiding future modeling of the RanBP2-type ZnF family. However, the mismatches in Zn-S distances and Zn-S-Cβ angles suggest there is still room for improvement in the current ZBM. These results suggest that folded domains act as structural modulators, indirectly modulating phase behavior by constraining IDR dynamics rather than engaging directly in multivalent contacts.

Utilizing the modified force field (ff99SBws-STQ-ZBM), our simulation of the condensed phase revealed interaction profiles that were remarkably consistent with those observed in single-chain ensembles. Despite the expected chain expansion and increased intermolecular contacts within the condensate, structured domains maintained their native folds while exhibiting minimal contributions to the overall interaction network. This conservation of interaction patterns between dilute and condensed phases aligns with previous observations in IDP systems,^31,54–56^ but extends these principles to multidomain proteins, demonstrating that folded domains primarily function as passive structural scaffolds rather than active participants in the multivalent interaction network driving phase separation. Quantitative contact analysis further revealed a nuanced interplay between interaction strength and residue abundance. While certain residues exhibit strong per-residue interactions, their overall contribution to LLPS may be limited by low abundance. Conversely, prevalent residues like glycine, despite forming weaker individual contacts, can significantly impact LLPS through their high frequency. This dual dependency framework, where less frequent residues with strong contributions coexist with high-abundance residues exerting substantial collective effects through numerous weak contacts, provides critical insight into the molecular determinants of phase separation in multidomain proteins. Notably, we identified FR π-π interactions – previously hypothesized but unverified – through sp²/π plane analysis.^2,31,87,98,103^ This multimodal interaction landscape illustrates how multidomain proteins leverage diverse physicochemical forces to fine-tune phase boundaries in complex systems.

In conclusion, our simulations provide detailed insights into the molecular mechanisms of FUS LLPS and its modulation by folded domain stability. While our modified force field successfully captured key aspects of FUS behavior – including folded domain stability and IDR interaction patterns – persistent discrepancies in global chain dimensions (e.g., R_g_ and R_h_) relative to experimental data highlight unresolved challenges in force field parameterization.

These limitations likely stem from an incomplete representation of solvent-mediated interactions and torsional potentials governing disordered regions.^51,52,104^ Beyond traditional interaction types, our analysis revealed significant FR contacts that participate in π-π stacking, expanding the repertoire of interactions contributing to FUS phase separation. However, the overall impact of these interactions on LLPS may be modulated by the low abundance of participating residues compared to more prevalent interactions. Future studies could address these gaps by integrating machine learning-derived corrections or experimental constraints from techniques like SAXS or single-molecule FRET.^105^ Importantly, our findings bridge a critical gap in the LLPS field by uncovering the interplay between folded and disordered regions in multidomain proteins. By demonstrating that single-chain simulations recapitulate condensate interaction profiles, we establish a computationally tractable framework for studying phase separation in other complex systems. These insights not only advance our mechanistic understanding of FUS-driven cellular organization but also provide a structural blueprint for designing therapeutics targeting specific interaction nodes – such as small molecules that stabilize RRM domains to inhibit pathological aggregation in ALS and FTD.

## Acknowledgements

We gratefully acknowledge support from NIGMS R35GM153388. We also thank the Texas A&M High Performance Research Computing (HPRC) for providing essential computational resources. The content of this work is solely the responsibility of the authors and does not necessarily represent the official views of the funding agencies.

