## Supporting information for "Elucidation of the molecular interaction network underlying full-length FUS conformational transitions and its phase separation using atomistic simulations"

**Elucidating the Molecular Interaction Network of Full-Length FUS via All-Atom Simulations**

| Force field | Detail | Protein | MD engine | Simulation time |
| --- | --- | --- | --- | --- |
| 03ws | ff03ws + CYZ | FL FUS | Amber | 10 μs * 3 replicas |
|  |  | RRM | Amber | 5 μs * 3 replicas |
|  |  | ZnF | Amber | 5 μs * 3 replicas |
| 99SBws-STQ | ff99SBws-STQ + CYZ | FL FUS | Amber | 5 μs * 3 replicas |
|  |  | RRM | Amber | 5 μs * 3 replicas |
|  |  | ZnF | Amber | 5 μs * 3 replicas |
| 03ws RRM restraint | ff03ws + CYZ + RRM restraints | FL FUS | OpenMM | 5 μs * 3 replicas |
| 03ws-ZBM | ff03ws + ZAFF | FL FUS | OpenMM | 5 μs * 3 replicas |
|  |  | ZnF | Amber | 5 μs * 1 replicas |
| 99SBws-STQ-ZBM | ff99SBws-STQ + ZAFF | FL FUS | OpenMM | 5 μs * 3 replicas |
|  |  | ZnF | Amber | 5 μs * 1 replicas |
| 03ws-ZBM RRM restraint | ff03ws + ZAFF + RRM restraints | FL FUS | OpenMM | 5 μs * 3 replicas |

**Supporting Table 1. Summary of the Performed Single-Chain All-Atom Simulations**

| Force field | ff99SBws-STQ + ZAFF, tip4p2005s |
| --- | --- |
| Number of chains | 25 |
| Number of amino acids per chain | 526 |
| Number of water molecules | 196161 |
| Number of Zn^2+^ | 25 |
| Number of Na^+^ | 556 |
| Number of Cl^-^ | 856 |
| Total number of atoms | 963231 |
| Equilibrated box dimension (nm) | 12.3 x 12.3 x 49.0 |
| Ionic strength (mM) | 158 |
| Amber 22 benchmark on A100 GPU | 52 ns / machine day |
| Amber 22 production | 2.5 μs NVT |

**Supporting Table 2. System Size of the Full-length FUS All-Atom Slab Simulation**

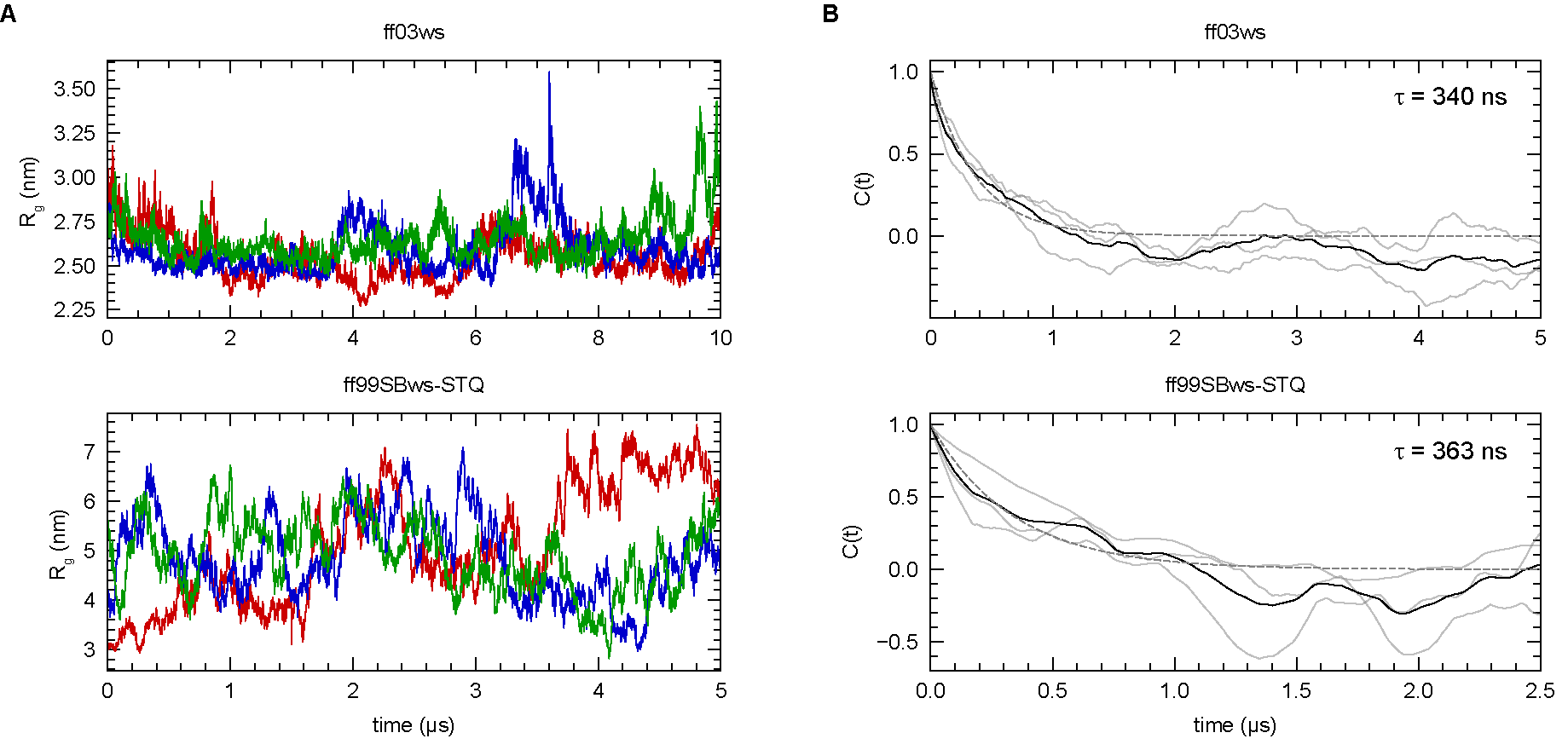

**Supporting Figure 1.** **Dynamic conformational ensemble in atomistic simulations. A.** R_g_ of FL FUS as a function of time from three independent replicas using two force fields with nonbonding parameters. **B.** R_g_ autocorrelation function (C(t)) for three independent replicas (grey) and mean correlation function (black). Conformational relaxation time is estimated by fitting the mean C(t) profile to a single exponential function (dashed black line).

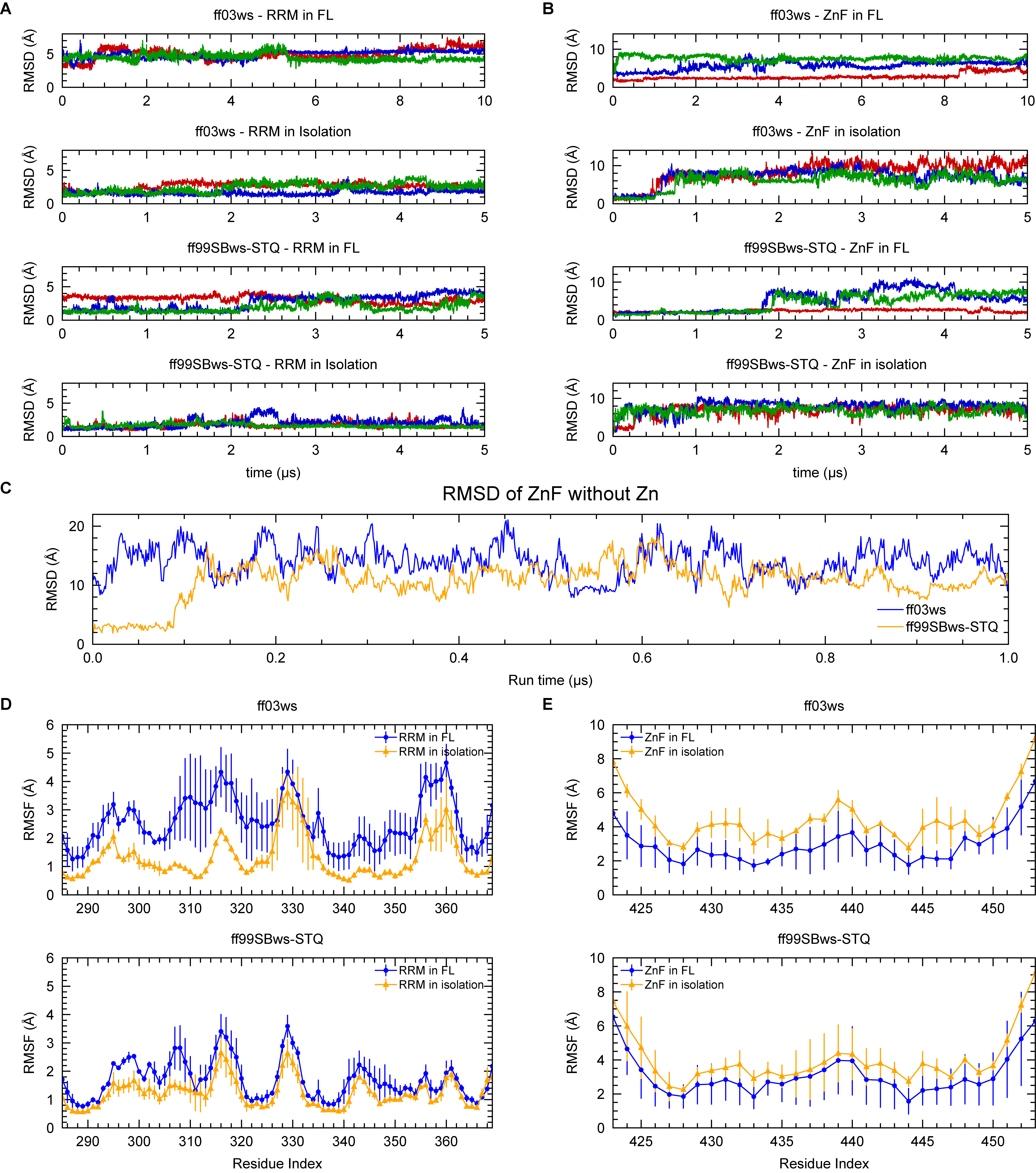

**Supporting Figure 2.** **Stability of folded domains.** **A. B.** Cα RMSD of the RRM and ZnF domains as a function of time across three independent replicas for each condition. **C.** Cα RMSD of the ZnF domain over time from isolated ZnF simulations without the zinc ion. **D. E.** Root-mean-square fluctuation (RMSF) of Cα atoms in the RRM and ZnF domains.

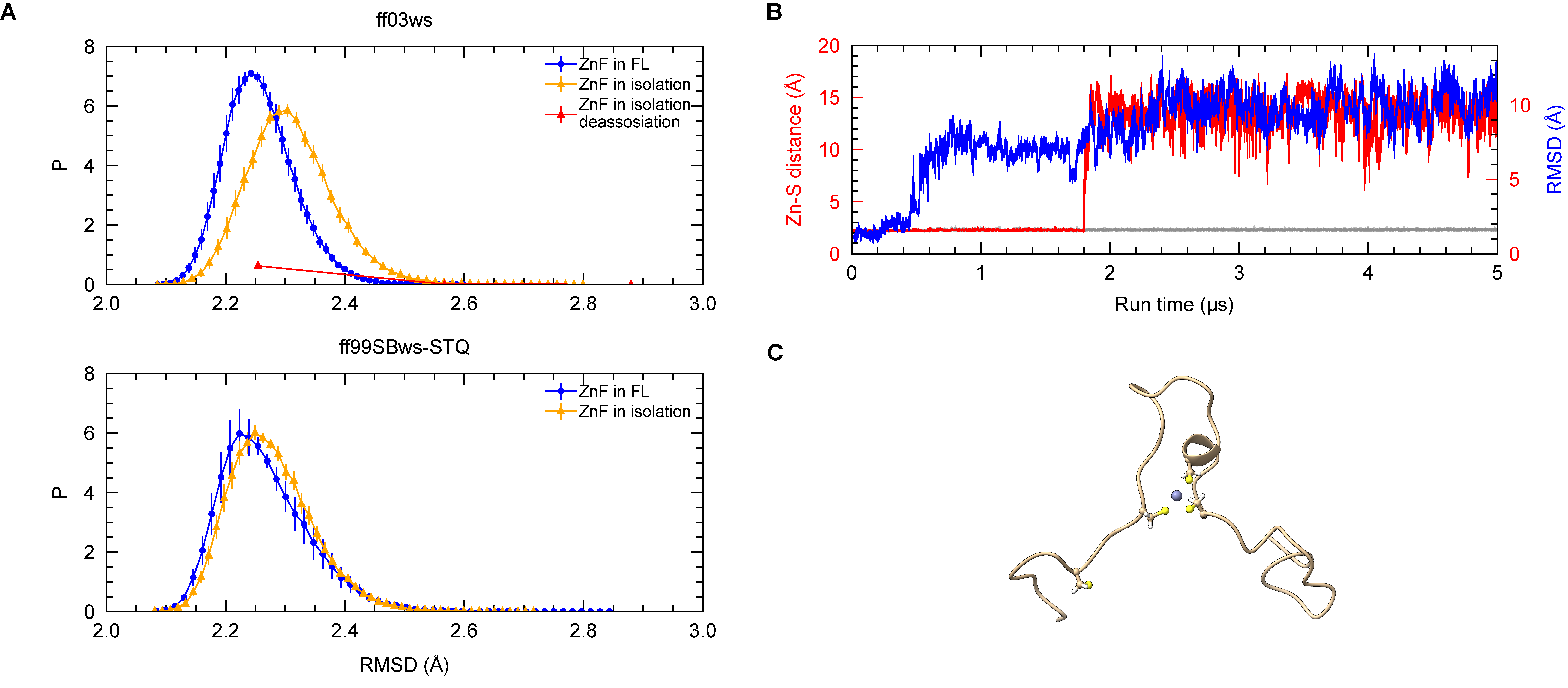

**Supporting Figure 3.** **Zinc-binding within the ZnF domain. A.** Distributions of Zn–S distances across all four cysteines for each condition from three replicas. One isolated ZnF simulation with ff03ws showed cysteine detachment. **B.** Zn–S distance (red) and RMSD (blue) over time for the detached cysteine instance. **C.** Snapshot of the detached ZnF structure. Cysteine residues and zinc were shown.

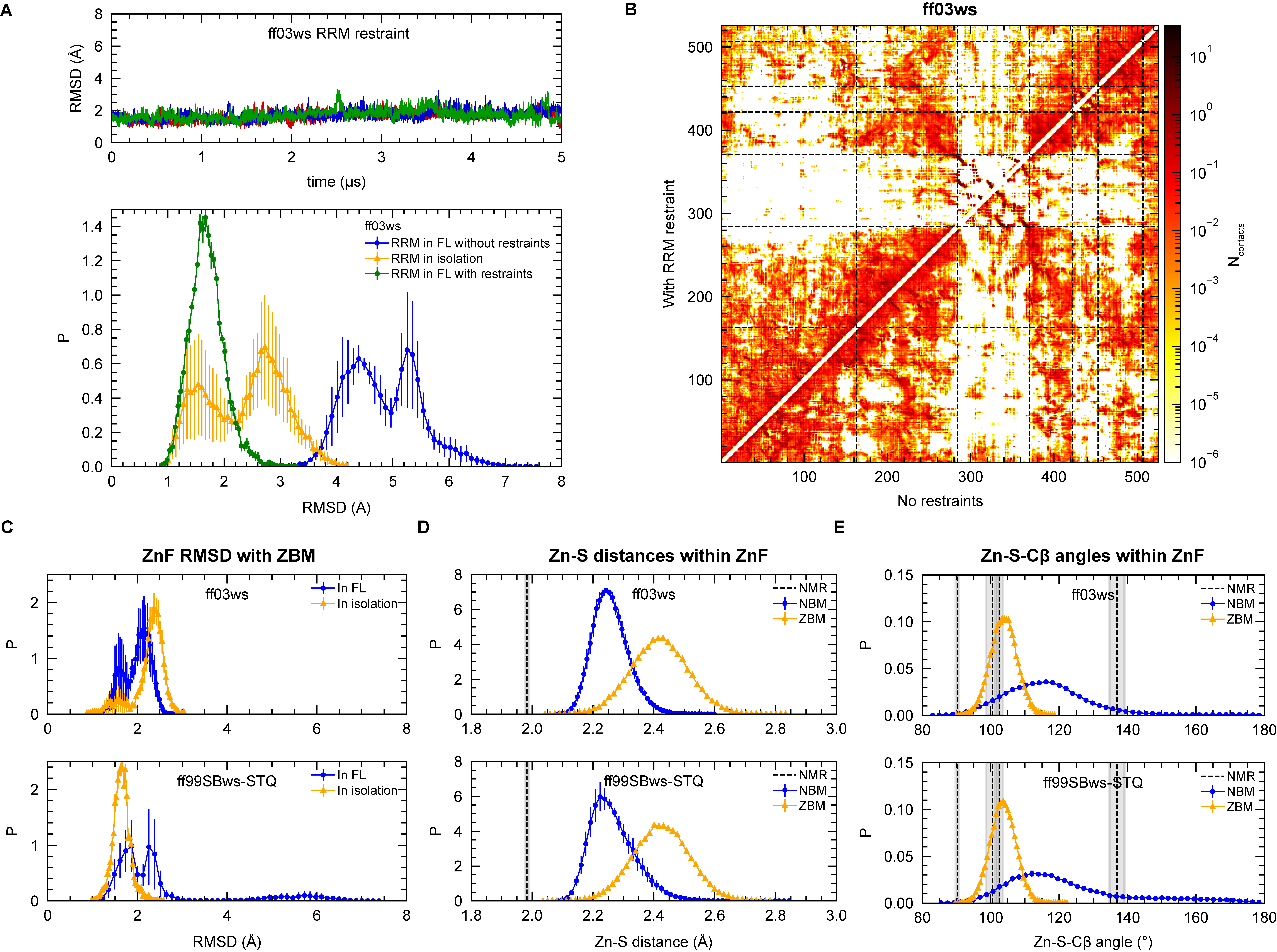

**Supporting Figure 4.** **Conformations between folded domains and full-length constructs. A.** Top: Cα RMSD of the RRM domain over time from three independent replicas of FL FUS simulations with ff03ws and RRM restraint. Bottom: Cα RMSD distributions of the RRM domain with ff03ws for FL FUS with and without RRM restraint, and for the isolated RRM without restraints. **B.** Intramolecular contact profiles from FL FUS single-chain simulations using ff03ws, with and without RRM restraints. **C.** Cα RMSD distributions of the ZnF domain using ZBM parameters, comparing FL FUS to the isolated ZnF domain. **D.** Distributions of Zn–S distances in the ZnF domain of FL FUS simulations using NBM or ZBM parameters, across three replicas and compared to the NMR ensemble. **E.** Distributions of Zn–S–Cβ angles in the ZnF domain of FL FUS simulations using NBM or ZBM parameters, across three replicas and compared to the NMR ensemble.

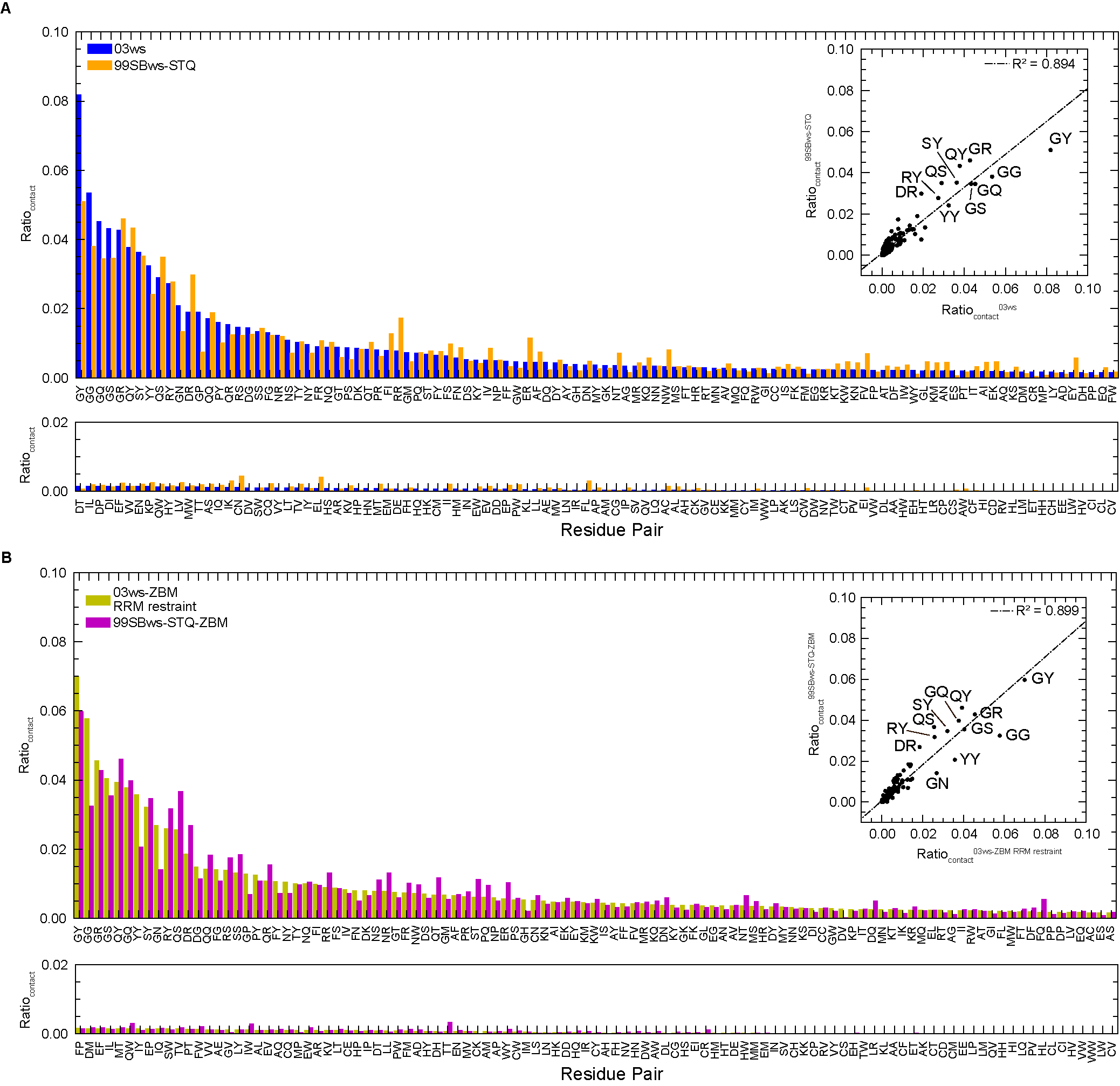

**Supporting Figure 5.** **Residue pair contacts from FL FUS single-chain simulations.** **A.** With unmodified force fields. Inset: Correlation between pairwise contact ratios in the ff03ws (x-axis) and ff99SBws-STQ (y-axis) models. **B.** With modified force fields. Inset: Correlation between pairwise contact ratios in the ff03ws-ZBM RRM restraint (x-axis) and ff99SBws-STQ-ZBM (y-axis) models. Same as **Figure 5** Inset.

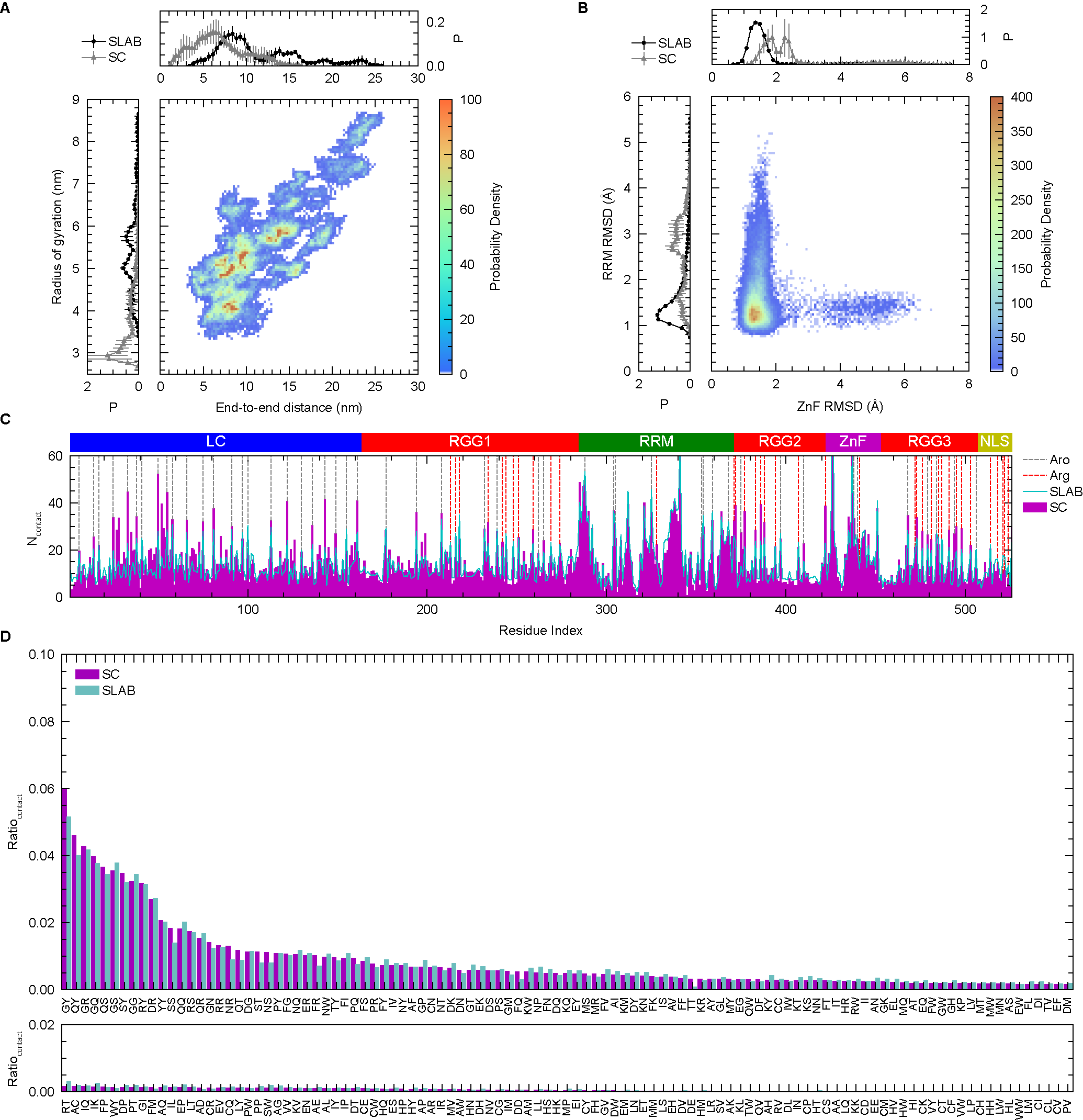

**Supporting Figure 6.** **FL FUS single-chain ensemble guides condensate formation.** **A.** Distribution of R_g_ and D_ee_ from FL FUS single-chain and slab simulations with ff99SBws-STQ-ZBM. **B.** Distribution of RMSD of the folded domains from FL FUS single-chain and slab simulations. **C.** One-dimensional summations of contacts from single-chain (SC) and condensate (SLAB) simulations. The positions of aromatic (Aro) and arginine (Arg) residues are indicated. **D.** Residue pair contacts from FL FUS slab and single-chain simulations.
